## Supplementary text for "Deep learning predictions of TCR-epitope interactions reveal epitope-specific chains in dual alpha T cells"

### Supplementary Information

Supplementary data - List of TCR-pMHCs sequence pairs (experimentally validated binders) used to train the MixTCRpred models

Supplementary Table 1 - List of all MixTCRpred models, number of TCRs binding to a specific pMHC, and AUC values for the 5-fold cross-validation test.

Supplementary Table 2 - HLAs of the donors in the 10X Genomics assay for scTCR sequencing of T-cells labeled with DNA-barcoded pMHC multimers

| | | | | |
|---|---|---|---|---|
| **Donor 1** | HLA-A*02:01 | HLA-A*11:01 | HLA-B*35:01 | - |
| **Donor 2** | HLA-A*02:01 | HLA-A*01:01 | HLA-B*08:01 | - |
| **Donor 3** | HLA-A*24:02 | HLA-A*29:02 | HLA-B*35:02 | HLA-B*44:03 |
| **Donor 4** | HLA-A*03:01 | HLA-A*03:01 | HLA-B*07:02 | HLA-B*57:01 |

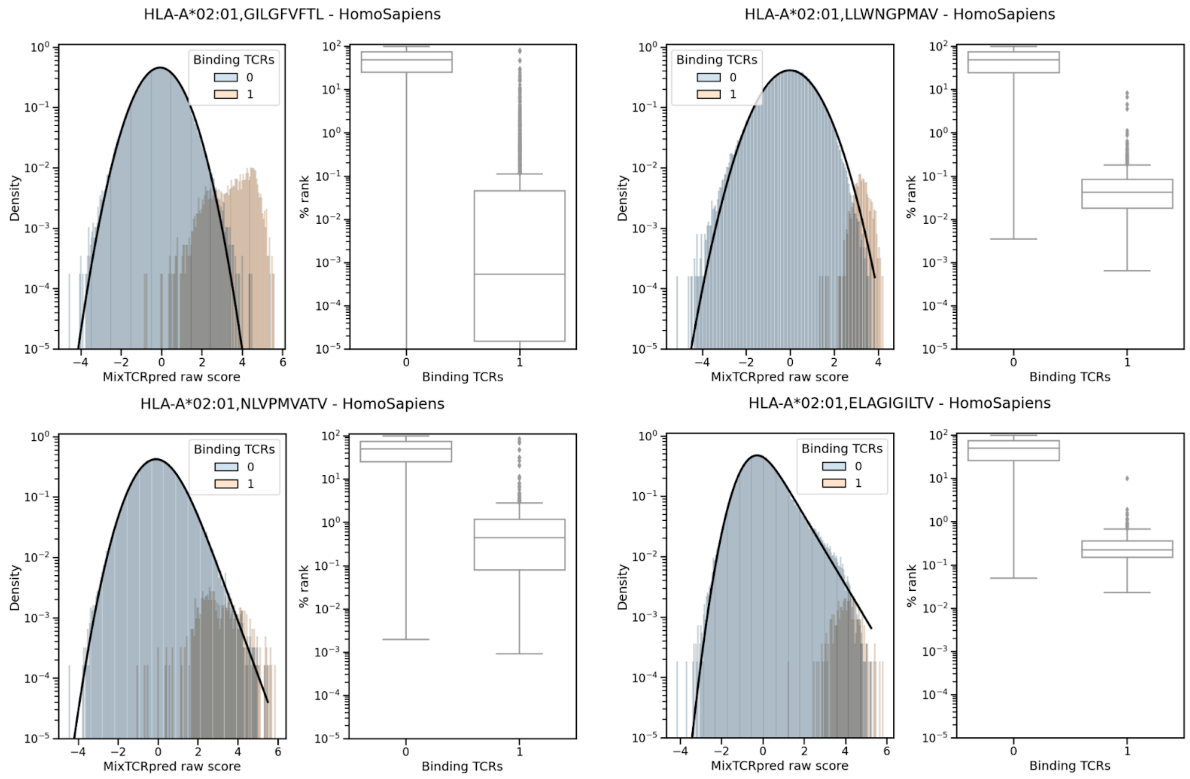


**Supplementary Figure 1.** Distribution of standardized MixTCRpred scores for $10^6$ random abTCRs with four epitopes. In black the fitted exponentially modified Gaussian distribution. The boxplots show the %rank for binding and non-binding TCRs.

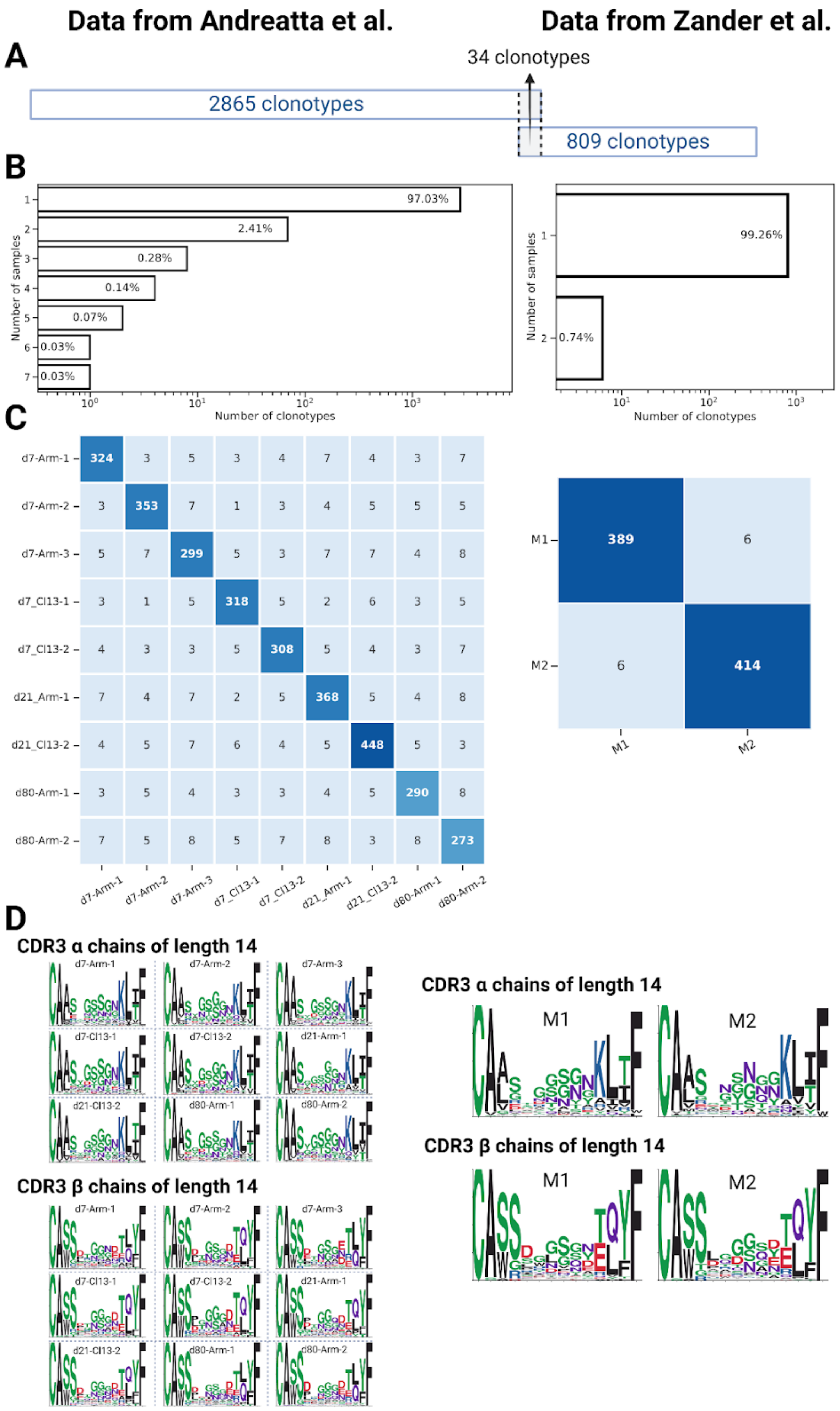
Data from Andreatta et al.
Data from Zander et al.
A
34 clonotypes
2865 clonotypes
809 clonotypes
B
97.03%
2.41%
0.28%
0.14%
0.07%
0.03%
0.03%
Number of samples
Number of clonotypes
99.26%
0.74%
C
d7-Arm-1
d7-Arm-2
d7-Arm-3
d7_Cl13-1
d7_Cl13-2
d21_Arm-1
d21_Cl13-2
d80-Arm-1
d80-Arm-2
M1
M2
D
CDR3 α chains of length 14
CDR3 β chains of length 14

**Supplementary Figure 2.** αβ TCRs from mice infected with the LCMV virus and specific for the H2-IAb, DIYKGVYQFKSV epitope. TCRs from the first study (left panels) were collected from 9 *Mus Musculus* samples infected with two different strains of LCMV (the Armstrong and Clone 13 strain), on day 7, 21 or 80 after infections [60]. In the second study (right panels) TCRs were collected from 2 samples infected with the LCMV-Clone 13 strain on day 10 after infection [61]. (A) Number of clonotypes from each study and the number of shared clonotypes. (B) Fraction of αβTCRs sequences that are unique to one sample (Number of samples=1, private clones) and shared between multiple mice (Number of samples > 1, public clones). (C) Number of αβTCRs shared between each pair of mice. (D) αβCDR3 sequences motifs for each sample separately. Only motifs for CDR3 regions of 14 amino acids are shown.

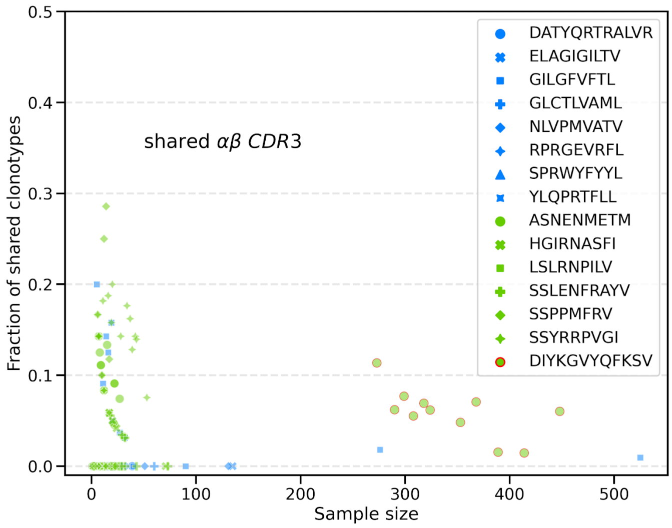


**Supplementary Figure 3.** Fraction of TCRs shared between the training (all samples but one) and the test set (the remaining sample) for the leave-one-sample-out cross-validation.

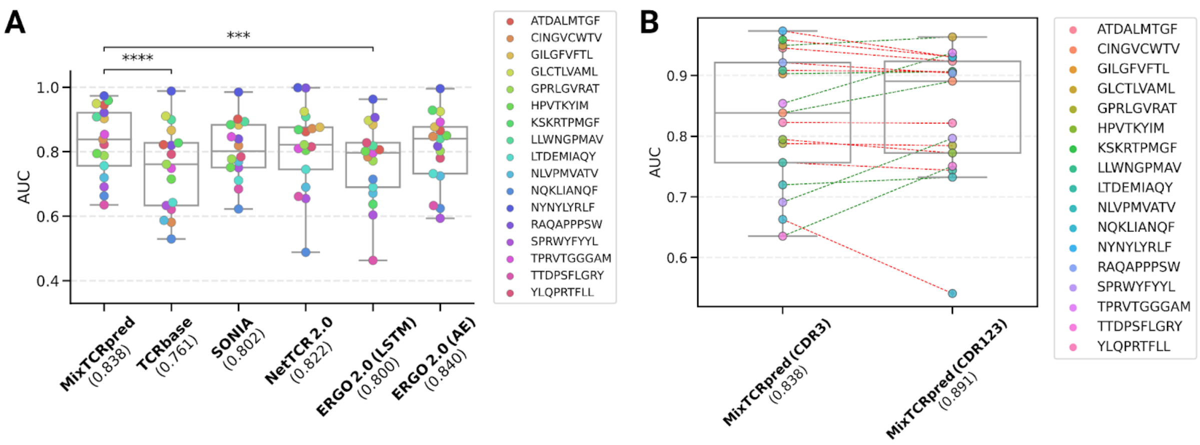


**Supplementary Figure 4.** (A) AUCs of CDR3-based tools tested on the benchmarking IMMREP22 dataset. (B) AUCs of MixTCRpred using as input features only the CDR3 sequences or the CDR1, CDR2 and CDR3 sequences.

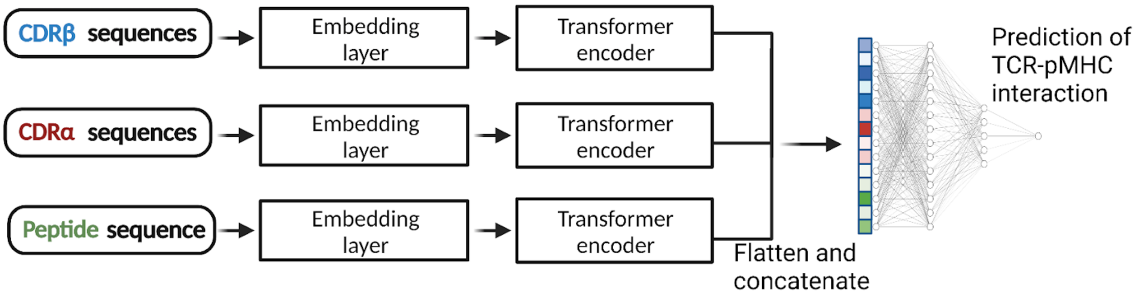


**Supplementary Figure 5**. Architecture of the pan-epitope MixTCRpred model. An additional encoding and transformer encoder layer was used for the peptide sequence.

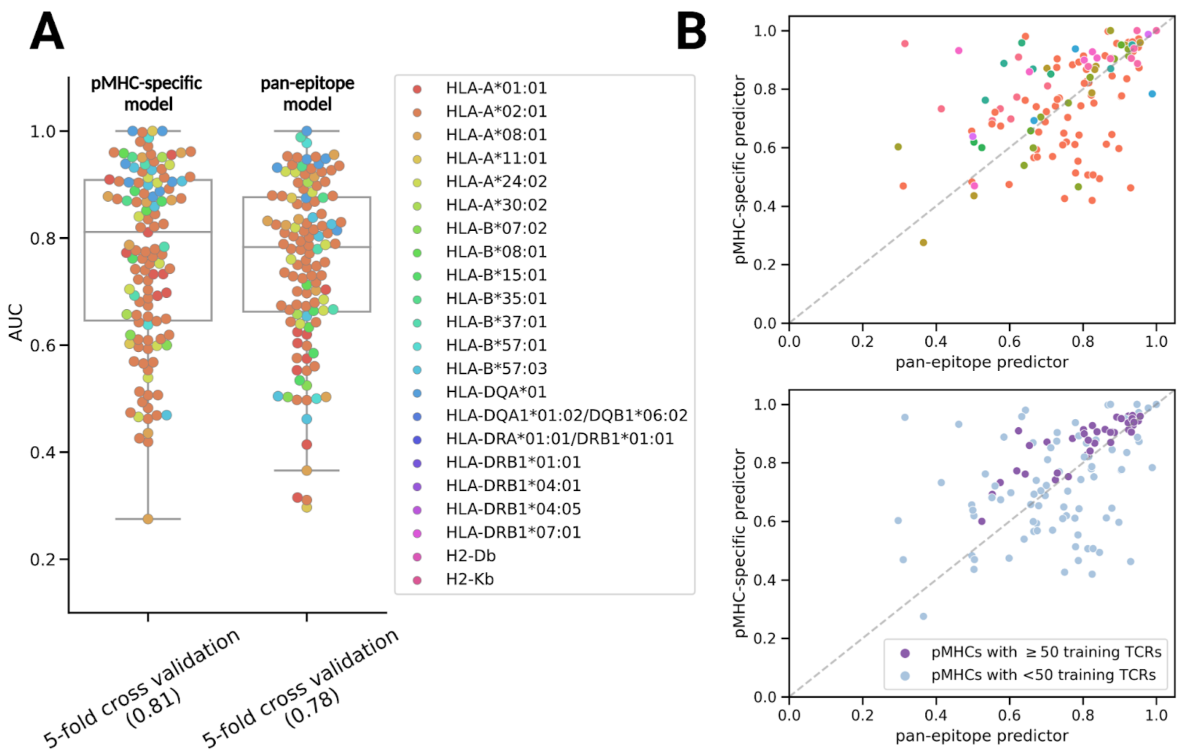


**Supplementary Figure 6.** (A) Comparison of AUCs of the pMHC-specific and the pan-epitope models for predicting TCRs interacting with epitopes already present in the training set. Each point is the average AUC 5-fold cross-validation, and different colors correspond to the shared MHC. (B) 5-fold cross-validation AUCs of the pan-epitope model vs. the AUCs of the pMHC-specific model, highlighting the shared MHCs (top plot) and if more than 50 TCRs are known for the pMHC (bottom plot).

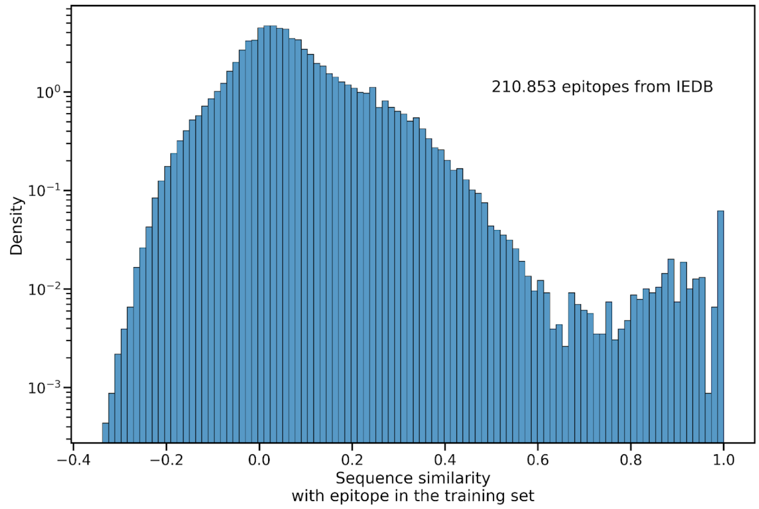

**Supplementary Figure 7.** Distribution of the sequence similarity between the 210.853 T cell epitopes from IEDB and the most similar epitope in the MixTCRpred training set.

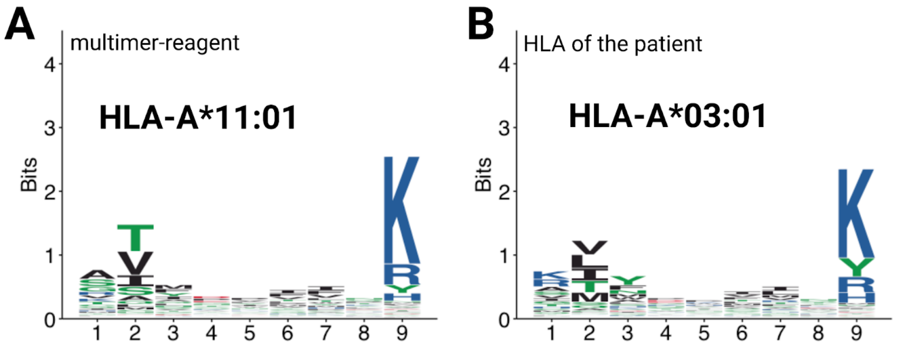


**Supplementary Figure 8.** Sequence motifs (downloaded from http://mhcmotifatlas.org/home [87]) of the HLA-A*11:01 (the HLA allele of the pMHC multimer reagent) and of the HLA-A*03:01 (HLA of the patient).

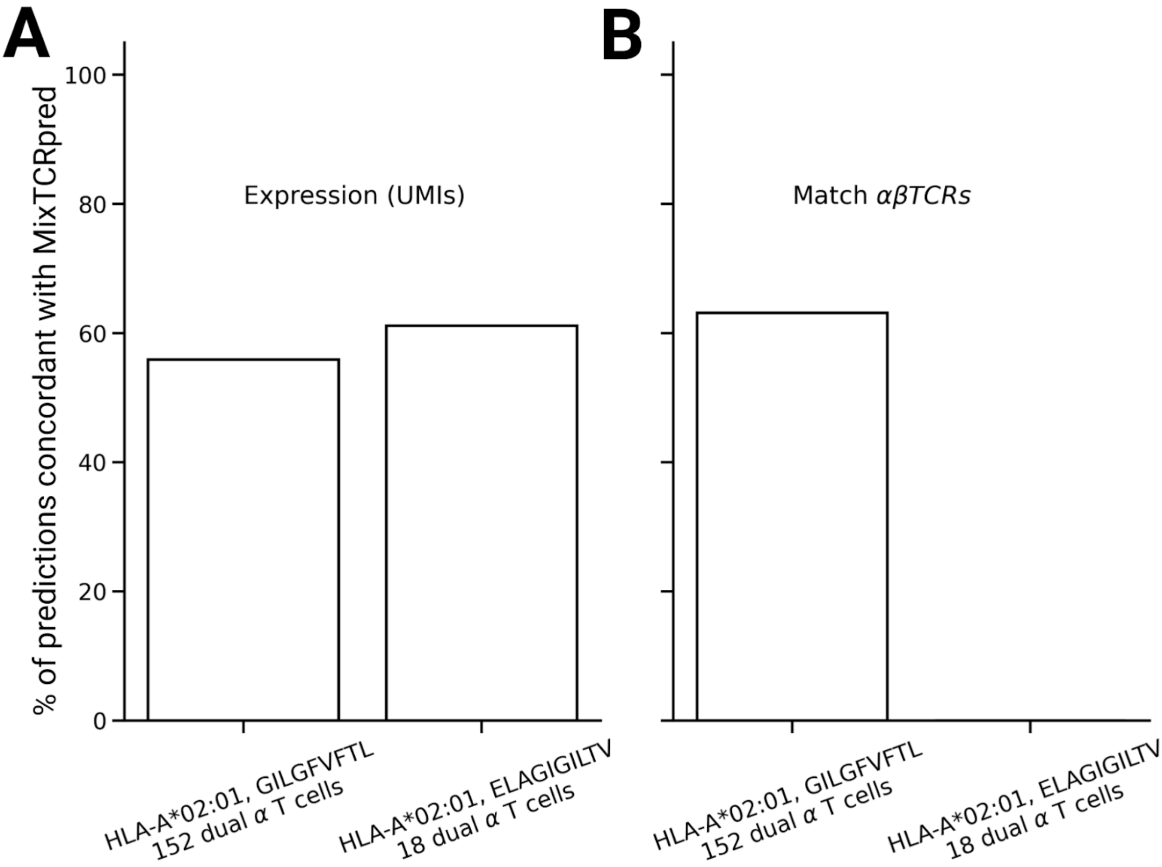

**Supplementary Figure 9.** Fraction of dual α T cells where the MixTCRpred predicted binder was also the more expressed α chain (A) or had an exact match in αβTCR sequence (B).

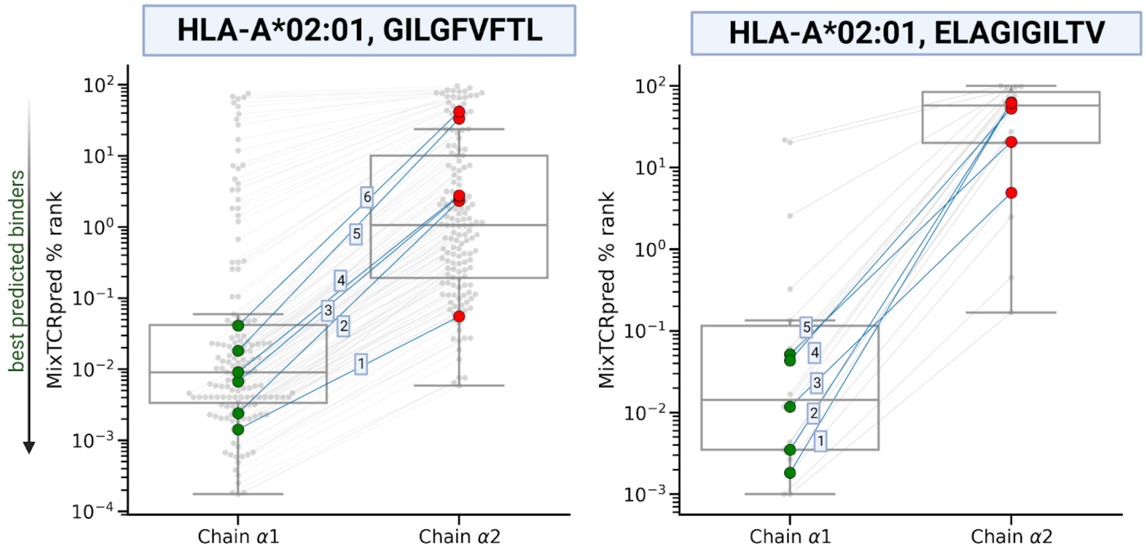


**Supplementary Figure 10.** MixTCRpred %ranks of TCRs from dual α T cells specific to HLA-A*02:01, GILGFVFTL and to HLA-A*02:01, ELAGIGILTV. Colored dots are TCRs used for experimental validation reported in Table 1. Green points are validated α chains, while red dots are non-binders.

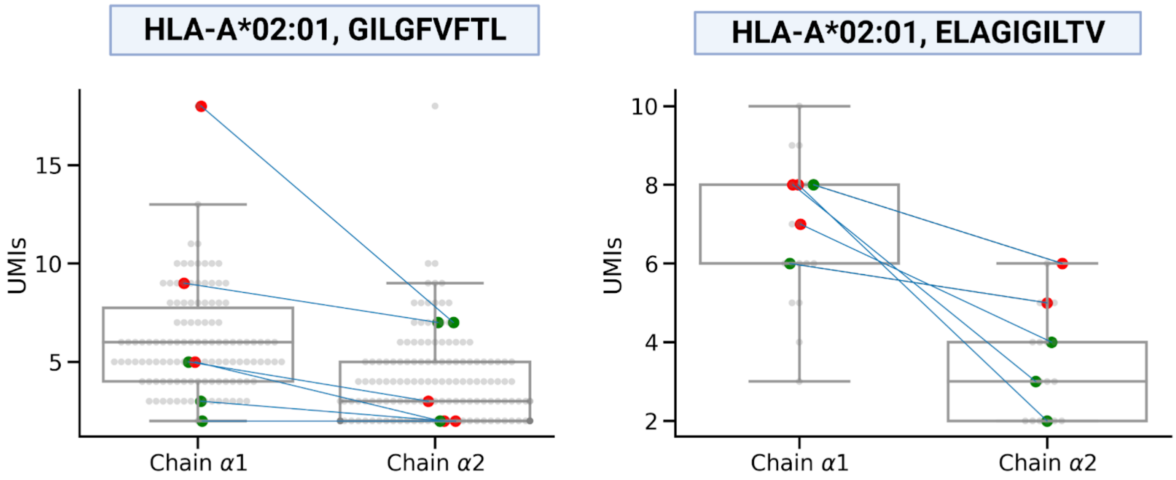


**Supplementary Figure 11.** UMI counts of TCRs from dual α T cells specific for HLA-A*02:01, GILGFVFTL and for HLA-A*02:01, ELAGIGILTV. In these plots chain α1 was defined as the chain with the higher level of expression. Colored dots correspond to the TCRs that were experimentally tested. The validated binders are depicted in green while the non-binders in red.

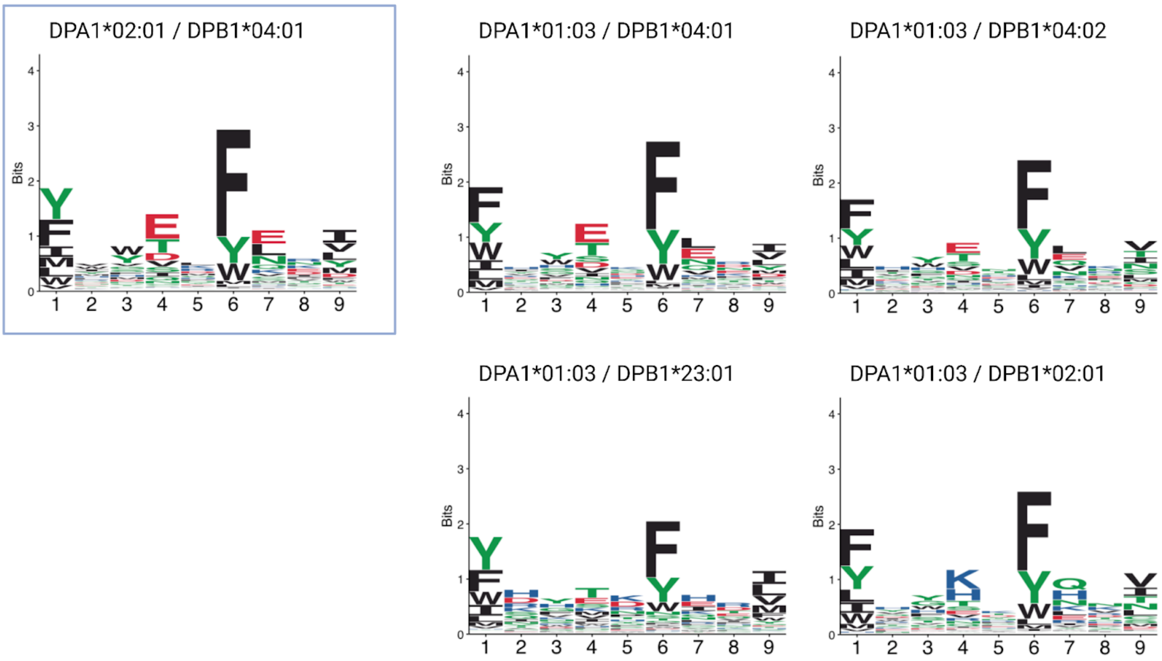


**Supplementary Figure 12.** Sequence motifs of DPA1*01:03 / DPB1*04:01 (i.e, the HLA allele to which the immunodominant SARS-CoV-2 TFEYVSQPFLMDLE epitope is restricted) and of other HLA-DP alleles.
